# Temporal expression of liver-stage malaria antigens shapes vaccine efficacy

**DOI:** 10.64898/2026.05.05.722570

**Authors:** Yu Cheng Chua, Lauren E. Holz, Daniel Fernandez-Ruiz, Sarah L. Draper, Regan J. Anderson, Benjamin J. Compton, Ralf B. Schittenhelm, Anton Cozijnsen, Charlie Jennison, Sophie Collier, Ryan W.J. Steel, Dhilshan Jayasinge, Stephanie Gras, Justin A. Boddey, Anthony W. Purcell, Geoff I. McFadden, Ian F. Hermans, Gavin F. Painter, William R. Heath

**Author notes:** These authors contributed equally to this work (Yu Cheng Chua, Lauren E. Holz). These authors contributed equally to this work (Ian F. Hermans, Gavin F. Painter, William R. Heath).

## Abstract

Vaccine-induced cytotoxic T cells can prevent malaria by killing parasite-infected hepatocytes during the liver stage. While several antigenic targets have been identified, little consideration has been given to their temporal expression. Here, we identified SERA1 of *Plasmodium berghei* as a late liver-stage target in rodent malaria and further showed that the classic vaccine antigen thrombospondin-related adhesion protein (TRAP) is only an early target. While vaccination with either antigen alone was modestly protective, combining these antigens enabled killing over the entire liver-stage, greatly improving efficacy. Given the relatively long liver-stage in human malaria, our findings imply TRAP-dependent vaccines likely utilize only a small proportion of the available liver-stage to eradicate parasites. Our findings further indicate that considerations of temporal coverage when selecting vaccine antigens will improve efficacy.

**One-Sentence Summary:** Temporally defining presentation of liver-stage antigens informs rational combinations that maximize malaria vaccine efficacy.

## Main Text

Malaria is a mosquito-transmitted disease caused by species of *Plasmodium* parasites. In 2022, this disease caused over 608,000 deaths, largely in African children (*1*). Plasmodium infections are complex: mosquito-derived sporozoites are released into the skin during a blood meal and migrate via the blood to the liver where they infect hepatocytes. Here they replicate extensively in a process that lasts 2 days in mice and about 7 days in humans. Merozoites then egress from hepatocytes to invade red blood cells in the blood-stage of infection, which triggers disease (malaria) and is responsible for all morbidity and mortality. As such, malaria can be prevented by killing parasites before they exit the liver.

While years of anti-parasite medication and insecticide-based interventions have substantially reduced deaths from malaria (*1*), such interventions are always under pressure from development of resistance to drugs and insecticides. Thus, a highly effective vaccine is greatly needed. Much progress has been made on this front, to the point that the RTS,S vaccine is now broadly approved for human use. However, this vaccine is only partially protective and immunity is short-lived (*2, 3*). R21, the newest licensed vaccine, necessitates three doses to achieve the WHO’s 75% efficacy target against clinical malaria in children, with an additional 12 month booster for sustained efficacy (*4-6*). This marks a significant milestone, though the pursuit of vaccines with greater efficacy is desirable, with several currently in development.

One vaccination strategy that is highly protective in naïve subjects (*7-9*), but has been difficult to extend to the field (*7, 10, 11*), is the administration of radiation attenuated sporozoites (RAS), which induce immunity during an aborted infection of the liver. RAS-induced protection is mediated by both antibodies that impair sporozoite migration and by CD8^+^ T cells that kill parasite-infected hepatocytes (*12*). More recently, liver-resident memory T cells (liver Trm cells) were shown to be the main subset of CD8^+^ T cells responsible for parasite killing (*13-16*). Liver Trm cells patrol sinusoids, scanning for infected hepatocytes (*14, 17, 18*). Several promising pre-clinical vaccination strategies have now been developed to favour liver Trm cell generation (*19, 20*), including our own (*14, 16, 21*). Development of subunit vaccines requires intricate knowledge of antigenic targets, several of which have been identified (*22*). However, most subunit vaccines have yielded relatively disappointing results in the field. Limited efficacy may be explained by two major problems. First, those vaccines tested in the field were developed before the discovery of Trm cells (*23, 24*) and generally focused on inducing relatively ineffective circulating T cell responses. Second, many antigens, including the two most popular targets, thrombospondin-related adhesion protein (TRAP) and circumsporozoite protein (CSP), show extensive strain polymorphisms (*25-28*), reducing specificity for field strains.

Recent development of vaccination strategies that efficiently generate liver Trm cells provided a solution to the first problem, though translation to humans awaits. To alleviate the second problem, new antigenic targets must be identified. As such, we recently used peptide elution and mass-spectroscopy to identify 60S ribosomal protein L6 (RPL6) as a protective antigen presented during *P. berghei* ANKA (PbA) infection of C57BL/6 (B6) mice (*29*). Importantly, its orthologue in the human pathogen *P. falciparum* is highly conserved (*29*). While continuing our search for new antigenic targets using this same approach, we identified SERA1 as a novel protective target. Although not a solution to the issue of polymorphism, its discovery revealed a third major problem associated with liver-stage vaccination. Here we show that variations in temporal expression can cause major limitations to vaccine efficacy and that these limitations extend to the canonical vaccine antigen, TRAP and potentially others.

## Results

### Discovery of SERA1 as an immunogenic liver-stage CD8^+^ T cell antigen

We previously described a mass spectroscopy-based approach to identify antigens naturally processed and presented on MHC I molecules by dendritic cells fed PbA-infected red blood cells (iRBC) (*29*). After screening previously infected and cured mice for immunity to a pool of 341 peptides 8-11 amino acids long, we identified a peptide from serine repeat antigen 1 (SERA1_1120-1130_) (PBANKA_0305100) as a potential antigenic target in mice vaccinated with either blood-stage parasites or with RAS (**Fig. 1A and B**). Compared to responses to epitopes from known proteins RPL6, TRAP and glideosome associated protein 50 (GAP50), the responses to SERA1 were weak, particularly in RAS vaccinated mice. Vaccination of mice lacking either H-2K^b^ or H-2D^b^ confirmed the reported restriction elements for RPL6 and GAP50 epitopes (*29, 30*) and suggested the SERA1 epitope was K^b^-restricted (**Fig. 1C**). Accordingly, a K^b^-tetramer was prepared to monitor SERA1-specific CD8^+^ T cell immunity by flow cytometry.

**Fig. 1.**
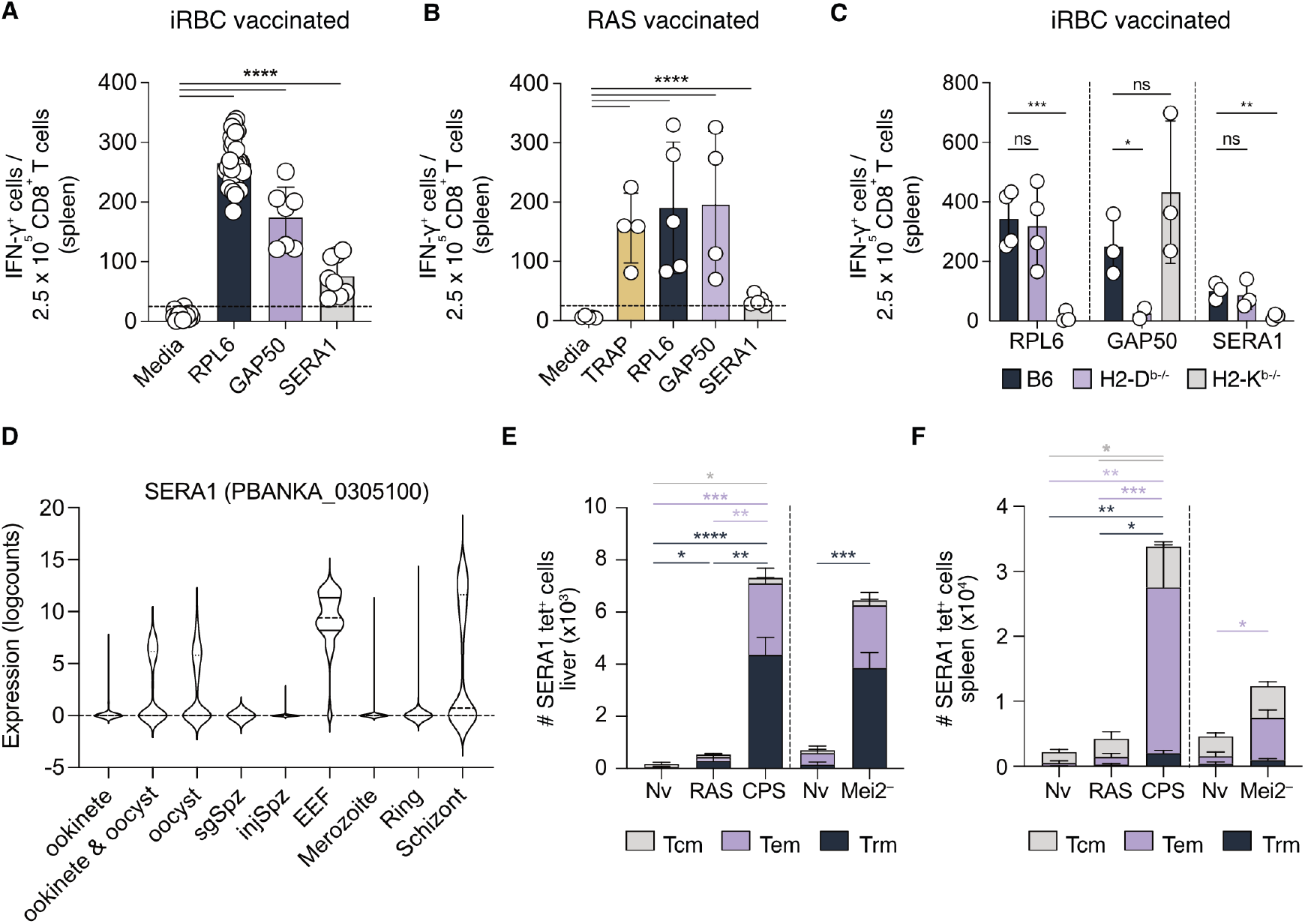
Identification of an immunogenic H2-K^b^-restricted SERA1 epitope expressed late during the liver stage. **(A)** Analysis of memory CD8^+^ T cell responses after iRBC immunization. B6 mice were infected i.v. with 10^6^ PbA iRBC and then cured of infection by treatment with chloroquine starting day 4 post-infection. 33 – 35 days later, 250,000 enriched CD8^+^ T cells from the pooled spleens of immunized mice were *in vitro* stimulated with or without peptide (media). 18-20 hours later, IFN-γ^+^ cells were enumerated by ELISpot assay. Data are pooled from >5 independent experiments. Each circle represents one experiment (n = 7-13 mice/experiment). **(B)** Analysis of memory CD8^+^ T cell responses after RAS immunization. B6 mice were immunized i.v. with 50,000 PbA RAS. 31 – 38 days later, 250,000 enriched CD8^+^ T cells from the pooled spleens of immunized mice were *in vitro* stimulated with or without peptide (media). 18-20 hours later, IFN-γ^+^ cells were enumerated by ELISpot assay. Data are pooled from 4-5 independent experiments. Each circle represents one experiment (n = 11-15 mice per experiment). **(C)** The SERA1-specific CD8^+^ T cell response is H-2K^b^ restricted. B6, H2-K^b-/-^ and H2-D^b-/-^ mice were immunized i.v. with 10^6^ PbA iRBC as described in (A). 32 – 39 days later, 250,000 enriched CD8^+^ T cells from the pooled spleens of immunized mice were *in vitro* stimulated with each of the indicated peptides.18 – 20 hours later, IFN-γ^+^ cells were enumerated by ELISpot assay. Data are pooled from 2 – 4 independent experiments. Each circle represents one experiment (n = 3-10 H2-K^b-/-^ mice/group; n = 3 – 11 H2-D^b-/-^ mice/group; n = 7 – 23 B6 mice/group, in each experiment). **(A-C)** Mean ± SD are shown. The dotted line in (A and B) represents the cut-off for a positive response calculated based on the mean spot count + 3SD of the media control. Data were log transformed and compared where indicated using the one-way ANOVA with Dunnett’s multiple comparison test. ns, not significant, \**P* < 0.05, \*\**P* < 0.01, \*\*\**P* < 0.001, \*\*\*\**P* < 0.0001. **(D)** Violin plots showing the expression of PbA *SERA1* (PBANKA_0305100) transcripts throughout the malaria life cycle. Data were adapted from publicly available single-cell transcriptomic data (*31*). Abbreviations: ookinete (bolus ookinete at 18 and 24 hours), ookinete & oocyst (48 hours from midgut tissue), oocyst (day 4 oocysts), sgSpz (salivary gland sporozoites), injSpz (sporozoites released by mosquito bite), EEF (liver stage), Ring (ring culture), schizont (late-stage cultures, including male, female, trophozoites and schizonts). **(E-F)** Analysis of memory CD8^+^ T cell responses after RAS, chemoprophylaxis with sporozoites (CPS) or *Mei2*^*–*^ sporozoite immunization. B6 mice were immunized twice (42-day interval) via the retro-orbital route with either 40,000 PbA RAS, or 40,000 wild-type PbA sporozoites followed by daily oral administration of chloroquine starting 4 hours post-infection for 7 days (CPS). 42 days after the last immunization, SERA1-specific memory CD8^+^ T cell responses were examined by flow cytometry. Tissue-resident memory (Trm) were identified as CD69^+^ CD62L^low^ cells, effector memory (Tem) were identified as CD69^-^ CD62L^low^ cells, and central memory (Tcm) were identified as CD69^-^ CD62L^+^ cells. Alternatively, B6 mice were immunized with three doses of 10,000 PbA *Mei2*^*–*^ sporozoites at 1-week intervals and then assessed for memory T cell responses 30 days later by flow cytometry. Total numbers of tet^+^ memory CD8^+^ T cells specific for SERA1 in the liver (E; individual data points (**fig. S5A**)) and spleen (F; individual data points (**fig. S5B**)). Data in (E and F) are pooled from two independent experiments (n = 5 – 12 mice/group). Mean ± SEM. Numbers for each memory T cell subset in (E and F) were log-transformed and compared using the one-way ANOVA with Tukey’s multiple comparison test or an unpaired two-tailed *t*-test. \**P* < 0.05, \*\**P* < 0.01, \*\*\**P* < 0.001, \*\*\*\**P* < 0.0001.

Examination of mRNA expression within the malaria cell atlas revealed expression of SERA1 in both the liver and blood stages of infection (*31, 32*) (**Fig. 1D)**. Western blot analysis of *P. berghi*-infected HepG2 cells suggests that SERA1 is only expressed towards the end of the liver-stage (*33, 34*). Late liver-stage expression was further supported by increased SERA1-specific CD8^+^ T cell immunity after vaccination by chemoprophylaxis with sporozoites (CPS), where live sporozoites are injected and allowed to progress through the liver stage unaffected, but then chloroquine administration is used to cure any emerging blood stage-infection (**Fig. 1E and F; fig. S1**). CPS contrasts vaccination with RAS, where the attenuated sporozoites have a truncated liver-stage. To exclude the possibility that enhanced immunity to SERA1 in CPS-vaccinated hosts was simply due to a response to the few blood stage parasites that briefly emerge before cure, we generated mutant parasites deficient for Meiosis inhibited 2 (*Mei2*)-like RNA binding protein (hereafter referred to as *Mei2*^*–*^), which causes aborted infection late in the liver stage (*35-37*). Strong SERA1-specific CD8^+^ T cell immunity in *Mei2*^*–*^ sporozoite-vaccinated mice further supported the late liver-stage expression of SERA1 (**Fig. 1E and F, right columns**).

### Immunization with SERA1 GLP vaccine generates abundant liver Trm cells and provides sterile protection against malaria

To test the protective capacity of SERA1_1120-1130_, we synthesised a glycolipid peptide (GLP) conjugate vaccine whereby this epitope was conjugated to a rearranged form of α-galactosylceramide using oxime ligation methodology as described previously (*21*). B6 mice were then immunized with either the SERA1 GLP, or a previously described RPL6 GLP as a positive control (*21*). Substantial numbers of memory CD8^+^ T cells were detected in the liver and spleen several weeks later (**Fig. 2A-D**), with significantly more SERA1-specific liver Trm cells than those specific for RPL6 (**Fig. 2B and C**), possibly due to the higher frequency of SERA1-specific naïve precursors (**Fig. 2E**).

**Fig. 2.**
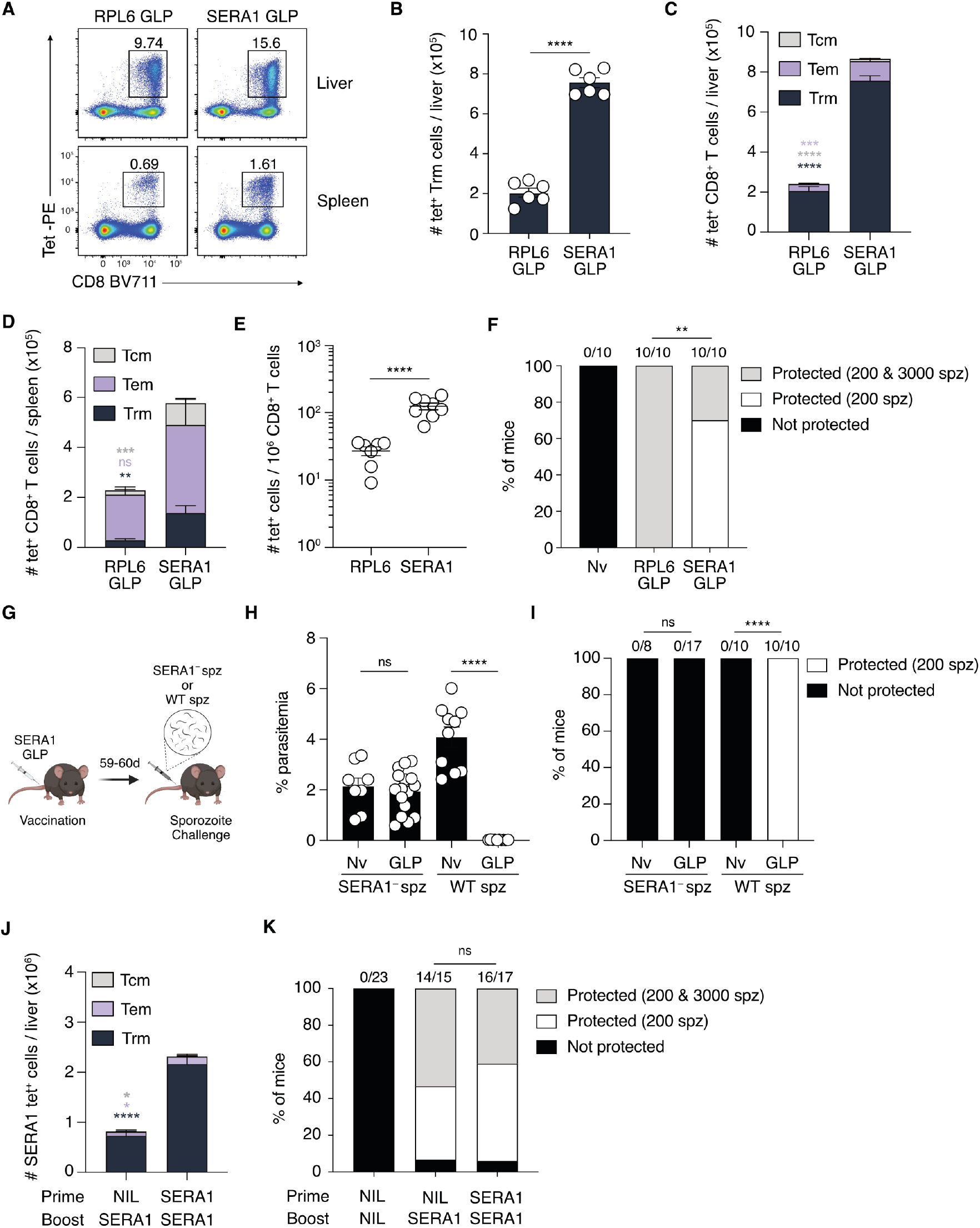
SERA1-specific liver Trm cells protect against PbA sporozoite challenge. **(A-D)** Analysis of memory CD8^+^ T cell responses after GLP vaccination. B6 mice were immunized intravenously with SERA1 GLP or RPL6 GLP vaccines and memory T cell responses were analyzed by flow cytometry 21-33 days later. **(A)** Representative FACS plots of CD8^+^ T cells specific for SERA1 or RPL6 in the liver (top row) and spleen (bottom row). **(B)** The numbers of tet^+^ CD8^+^ liver Trm cells specific for SERA1 and RPL6 in individual mice (circles). Mean ± SEM. **(C and D)** The total numbers of tet^+^ memory CD8^+^ T cells specific for SERA1 and RPL6 in the liver (C; individual data points (**fig. S6A**)) and spleen (D; individual data points (**fig. S6B**)). Data in (A-D) are pooled from two independent experiments (n = 6 mice/group). Data in (B) were log-transformed and compared using an unpaired two-tailed *t*-test. Numbers for each memory T cell subset in (C and D) were log-transformed and compared using an unpaired two-tailed *t*-test. ns, not significant, \*\**P* < 0.01, \*\*\**P* < 0.001, \*\*\*\**P* < 0.0001. **(E)** Number of tet^+^ CD8^+^ T cells per 10^6^ naïve CD8^+^ T cells in individual mice (circles). Mean ± SEM. Data are pooled from three independent experiments (n = 7 – 8 mice/group). Data were log-transformed and compared using an unpaired two-tailed *t*-test. \*\*\*\**P* < 0.0001. **(F)** SERA1 GLP and RPL6 GLP-vaccinated mice were challenged intravenously with 200 PbA sporozoites >27 days after vaccination, followed by rechallenge of surviving mice with 3000 PbA sporozoites intravenously at day 110 post-vaccination. After each challenge, sterile protection was assessed by measuring parasitemia at different time points up to day 12 post-infection. Shown is the proportion of mice that succumbed to the 200 sporozoite challenge (black bars), were protected against the 200 sporozoite challenge only (white bars) or were protected against both the 200 and 3000 sporozoite challenges (grey bars). Numbers above bars indicate the number of mice that were protected against the 200 sporozoite challenge over the total number of mice challenged. Data are pooled from 2 independent experiments (n = 10 mice/group). For 3000 sporozoite challenges, the naïve group is not shown, but all mice succumbed to malaria infection. Percent sterile protection after the 3000 sporozoite challenge was compared using a two-sided Fisher’s exact test. \*\**P* < 0.01. **(G)** B6 mice were immunized intravenously with SERA1 GLP and then challenged with 200 PbA wild-type (WT) or *SERA1*^*-*^ sporozoites 59 – 60 days later. **(H)** The percentage of parasitemia at day 7 post-infection. Mean ± SEM. **(I)** The proportion of mice that succumbed to malaria infection (black bar) or were protected against the 200 sporozoite challenge (white bar). Numbers above bars indicate the number of mice that were protected against the 200 sporozoite challenge over the total number of mice challenged. Data are pooled from 2 independent experiments (n = 8 -17 mice/group). Data in (H) were log-transformed and compared using an unpaired two-tailed *t*-test. Percent sterile protection after the 200 sporozoite challenge in (I) was compared using a two-sided Fisher’s exact test. ns, not significant, \*\*\*\**P* < 0.0001. **(J-K)** Analysis of the SERA1-specific memory CD8^+^ T cell response after prime-boost immunization with SERA1 GLP (60-day interval). **(J)** Total number of SERA1 tet^+^ memory CD8^+^ T cells in the liver at 32 – 53 days after the last immunization (individual data points (**fig. S6C**)). Mean ± SEM. Data are pooled from 3 independent experiments (n = 11 – 14 mice/group). Numbers for each memory T cell subset were log-transformed and compared using an unpaired two-tailed *t*-test. \**P* < 0.05, \*\*\*\**P* < 0.0001. **(K)** Sterile protection against sporozoite challenge. Separate cohorts of vaccinated mice were challenged intravenously with 200 PbA sporozoites and sterile protection was assessed by measuring parasitemia up to day 12 post-infection. On day 138 – 155 post-vaccination, surviving mice from the 200 sporozoite challenge were injected intravenously with 3000 PbA sporozoites and sterile protection again assessed. Shown is the proportion of mice that succumbed to the 200 sporozoite challenge (black bars), were protected against the 200 sporozoite challenge only (white bars) or were protected against both the 200 and 3000 sporozoite challenges (grey bars). Numbers above bars indicate the number of mice that were protected against the 200 sporozoite challenge over the total number of mice challenged. Data are pooled from 4 independent experiments (n = 15 – 23 mice/group). For 3000 sporozoite challenges, the naïve group is not shown, but all mice succumbed to malaria infection. Percent sterile protection after the 3000 sporozoite challenge was compared using a two-sided Fisher’s exact test. ns, not significant.

To assess the protective efficacy of the SERA1 GLP vaccine, immunized mice were challenged i.v. with a small but lethal dose of 200 PbA sporozoites (equivalent to about 1 mosquito bite (*38*)), and then monitored for blood-stage parasitemia for 12 days. This revealed 100% sterile protection by SERA1 GLP, as also seen in the positive control group of RPL6 GLP vaccinated mice (**Fig. 2F**). Surviving mice were then rechallenged with a higher dose of 3000 PbA sporozoites, which showed that while some SERA1-vaccinated mice were protected from this dose, such protection was significantly less than measured for RPL6-vaccinated mice, where all mice were protected (**Fig. 2F**). Of note, the observed protection was liver-stage specific because similar SERA1-immunized mice succumbed to blood-stage infection when challenged with 500 PbA iRBC (**fig. S2A and B**). Protection was also shown to be SERA1-specific as mice challenged with SERA1-deficient parasites were not protected (**Fig. 2G-I**). Finally, while a booster vaccination with SERA1 GLP increased liver Trm cell numbers significantly (**Fig. 2J**), it still failed to protect all mice from a 3000 sporozoite challenge (**Fig. 2K**). Together, these findings showed that SERA1 was an excellent antigenic target for recognition by liver Trm cells, though its late liver-stage expression may have hampered complete eradication of large numbers of parasites.

### TRAP is a suboptimal target for liver Trm cells but can enhance immunity when combined with SERA1

TRAP mRNA is predominantly expressed by sporozoites, with expression dramatically decreasing upon liver infection (*32, 39, 40*). As such, most MHC I-TRAP complexes presented by infected hepatocytes are likely derived from TRAP protein carried into the hepatocyte by the sporozoite. These complexes will have a limited (but unknown) period of presentation on the hepatocyte surface before being lost to natural MHC I-peptide turnover (*41*) at which time host cells will no longer be susceptible to killing by TRAP-specific T cells. We have extensively examined the liver Trm cell response to TRAP and found that more Trm cells are required to achieve sterile protection against TRAP than are required for RPL6 (*29*), the latter of which is expressed throughout the liver stage (*31, 32*). One possible explanation for this difference is that TRAP is only available as an early liver-stage target. With this concept in mind and knowledge that SERA1 is a late liver-stage antigen, we hypothesized that combining liver Trm cell responses to each of these antigens would augment liver-stage immunity by providing killing capacity at both early and later stages of infection. To test this hypothesis, groups of B6 mice were prime-boost vaccinated with either TRAP GLP or SERA1 GLP or a combination of both. To prevent the total number of Trm cells in the combined vaccination group from exceeding the total number in the SERA1 alone group, SERA1 was not used to boost the combined group. This circumvents any improvement in protection in the combined group from being attributed to greater numbers of total liver Trm cells and enables gains to be attributed to dual antigen specificity. Examination of memory responses on day 60 showed that these vaccination approaches induced ∼50,000 TRAP Trm cells and ∼1,500,000 SERA1 Trm cells (**Fig. 3A, B and fig. S3**), with the total number of Trm cells in the SERA1 GLP prime-boost group similar to but trending higher than the total combined number of Trm cells of both specificities in the SERA1 GLP + TRAP GLP group (**Fig. 3B**). When these groups were challenged with 3000 sporozoites, TRAP vaccination alone showed almost no sterile protection (**Fig. 3C**), but the relatively low parasitemia detected on day 6 compared to the unvaccinated group (**Fig. 3D**) indicated substantial killing of infected hepatocytes. Vaccination with SERA1 alone showed the usual suboptimal sterile protection with 10 of 15 mice protected (**Fig. 3C**). In contrast, mice vaccinated with both antigens were fully protected. As there was no significant difference in the number of liver Trm between this group and the SERA1 GLP alone group (**Fig. 3B**), we concluded that the combined action of TRAP and SERA1 Trm cells was more effective than a similar number of SERA1 Trm cells acting alone. One way to explain this finding is if TRAP Trm cells killed early in the liver stage and then SERA1 Trm cells killed late, with their combined activity providing improved efficacy.

**Fig. 3.**
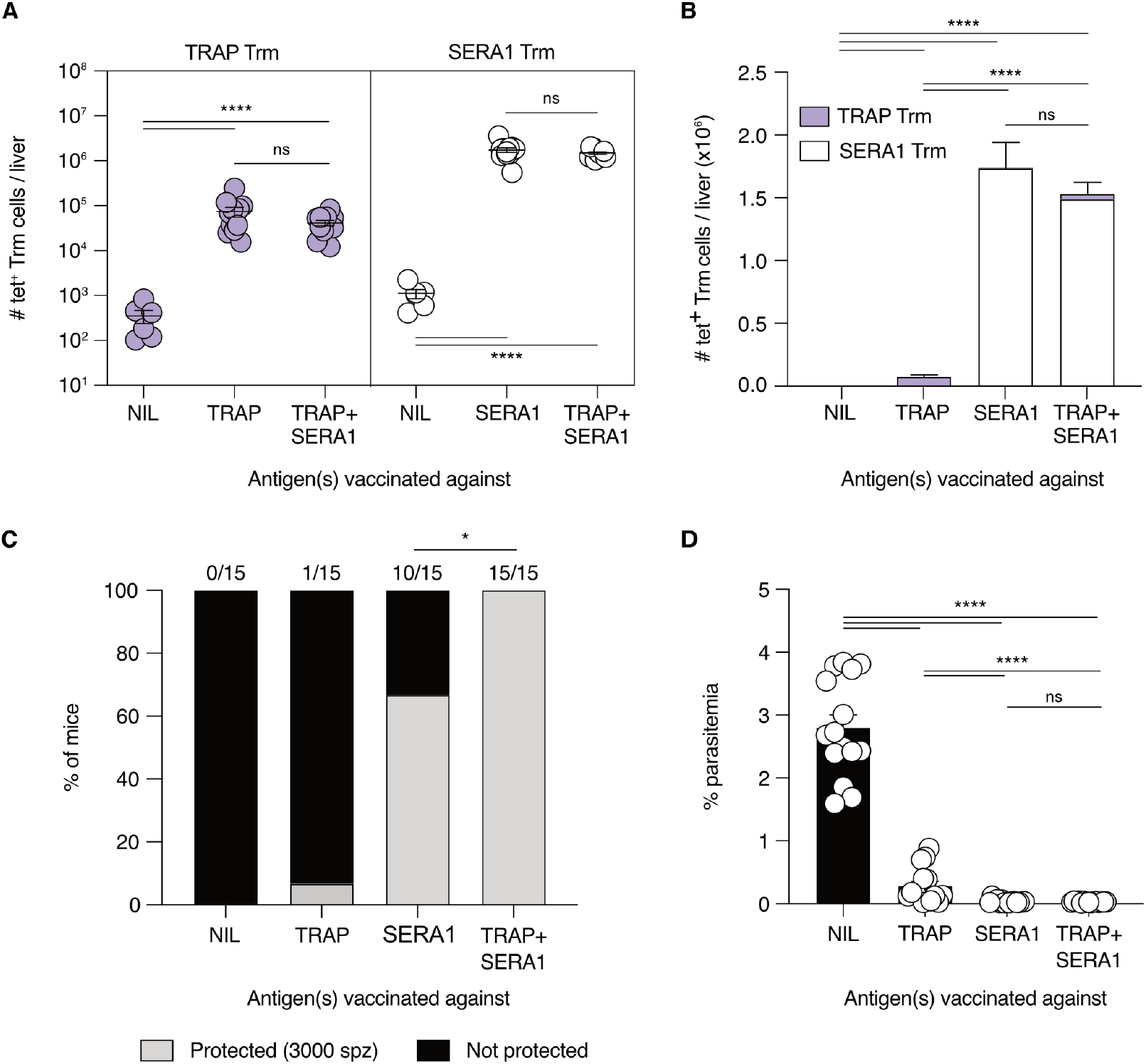
Combining liver Trm cell responses to SERA1 and TRAP improves protection. (**A and B**) Liver Trm responses. SERA1 and TRAP CD8^+^ Trm cell responses after vaccination with single or combined antigens. B6 mice were left unvaccinated (NIL), or primed and boosted intravenously with SERA1 GLP (SERA1) or TRAP GLP (TRAP) vaccines. Alternatively, B6 mice were immunized intravenously with an admix of SERA1 GLP and TRAP GLP vaccines, followed by an intravenous booster with TRAP GLP vaccine (TRAP+SERA1). Memory CD8^+^ T cell responses were analyzed by flow cytometry 30 days after the last vaccination (see fig. S3 for representative FACS plots). **(A)** The numbers of tet^+^ CD8^+^ liver Trm cells specific for TRAP (purple) or SERA1 (white) in individual mice (circles). Mean ± SEM. Data are pooled from 3 independent experiments (n = 6 – 12 mice/group). Data were log-transformed and compared using the one-way ANOVA with Tukey’s multiple comparison test. ns, not significant, \*\*\*\**P* < 0.0001. **(B)** The total number of PbA-specific CD8^+^ Trm cells per mouse in each vaccination group. Mean + SEM. The average number of each specificity is highlighted with TRAP (purple) and SERA1 (white). Data were log-transformed and compared using the one-way ANOVA with Tukey’s multiple comparison test. ns, not significant, \*\*\*\**P* < 0.0001. **(C and D)** Protection against sporozoite challenge. Separate cohorts of vaccinated mice were challenged intravenously with 3000 PbA sporozoites and sterile protection was assessed by the lack of parasitemia at different time points up to day 12 post-infection. (**C**) The proportion of mice that succumbed to malaria infection (black bars) or were protected against the 3000 sporozoite challenge (grey bars). Numbers above bars indicate the number of mice that were protected against 3000 sporozoite challenge over the total number of mice challenged. (**D**) Day 6 parasitemia. Mean ± SEM. Data in (C and D) are pooled from 3 independent experiments (n =15 mice/group). Data in (C) was compared using a two-sided Fisher’s exact test. \**P* < 0.05. Data in (D) were log-transformed and compared using the one-way ANOVA with Tukey’s multiple comparison test. ns, not significant, \*\*\*\**P* < 0.0001.

### Liver-stage parasites are killed early by TRAP Trm cells and late by SERA1 Trm cells

The above finding supported the idea that protection against TRAP and SERA1 operated in different temporal windows. To directly examine the kinetics of liver-stage parasite killing by these Trm cells, we used luciferase-expressing parasites to monitor killing at different times. Mice were vaccinated as previously (**Fig. 3**) against TRAP, SERA1 or both antigens, and then 30 days later each group was challenged with 30,000 luciferase-expressing PbA sporozoites (*42*). Luciferase activity was then measured in the liver at 24 and 42 hours to assess parasite growth (**Fig. 4A and B**). By comparing parasite growth in unvaccinated vs vaccinated mice, we could assess parasite killing to each antigen or the combination. The significant reduction in parasite load in the first 24 hours for mice immunized to TRAP compared to unvaccinated mice (NIL group) showed that killing to this antigen occurred early in the liver stage (but did not exclude additional late killing). In contrast, the equivalent parasite loads detected in SERA1 vaccinated mice and unvaccinated mice at 24 hours indicated a lack of killing to SERA1 early. At 42 hours, however, SERA1-vaccinated mice showed a significant reduction in parasite load compared to unvaccinated mice, indicating killing to this antigen occurred between 24 and 42 hours. The significant reduction in parasite load at 42 hours for mice vaccinated with a combination of both antigens, compared to mice vaccinated with either antigen alone, showed that by combining these antigens more parasites could be killed.

**Fig. 4.**
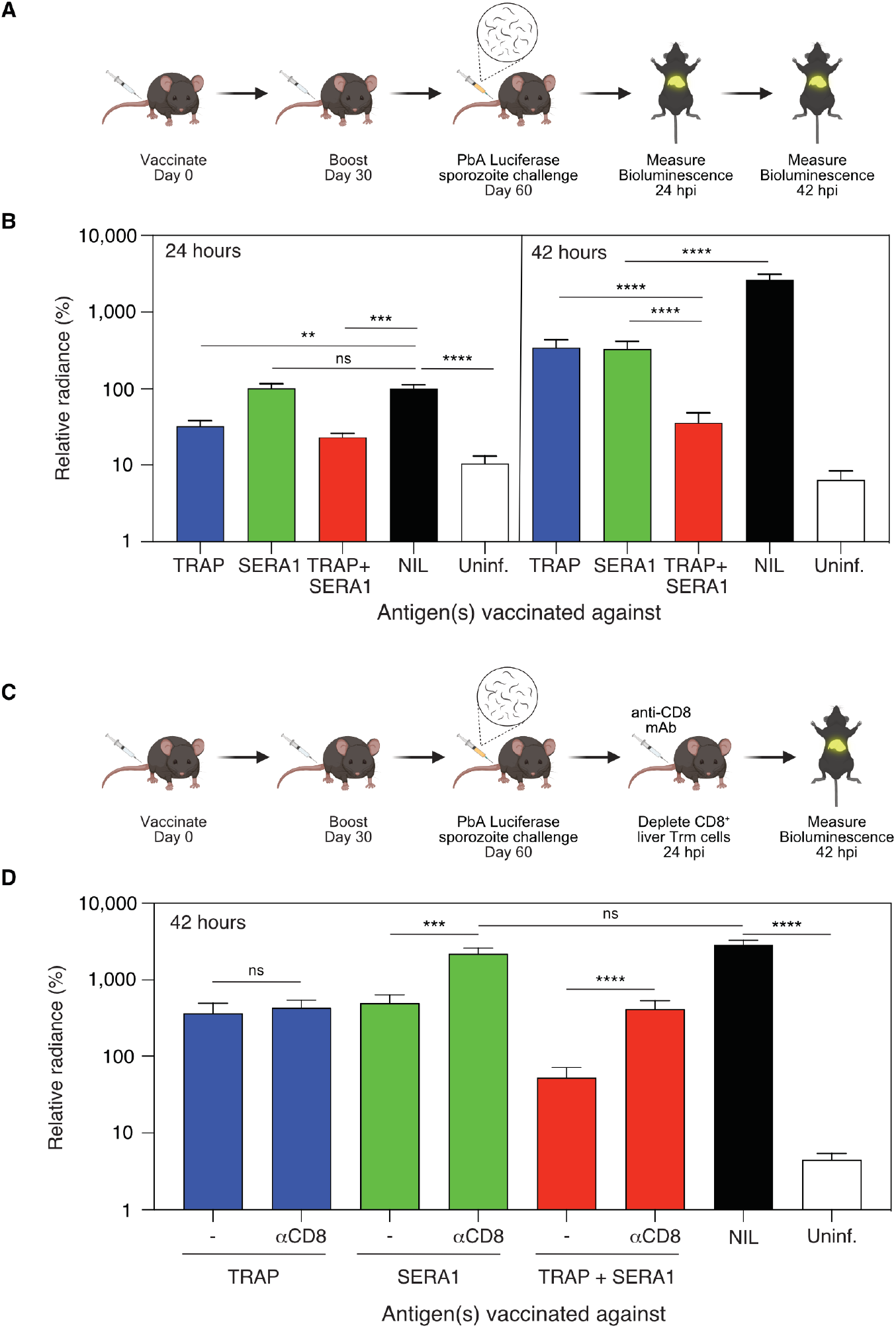
Vaccination against SERA1 and TRAP provide late and early liver-stage parasite control, respectively. **(A)** SERA1 only provides late liver-stage control. Schematic for (B). **(B)** A group of B6 mice were immunized intravenously with an admix of SERA1 GLP and TRAP GLP vaccines, followed by an intravenous booster with TRAP GLP vaccine (TRAP+SERA1) 30 days later. Alternatively, other groups of B6 mice were primed and boosted intravenously with SERA1 GLP (SERA1) or TRAP GLP vaccines (TRAP). One-month post-boost, mice were challenged with 30,000 luciferase expressing PbA sporozoites and imaged by IVIS at 24 (left) and 42 hours (right). To normalize data, % relative radiance is shown. This value represents the percent bioluminescence flux for each mouse relative to the average bioluminescence for the 24-hour readings in the unvaccinated (NIL) group. Mean ± SEM (n = 17 – 36 mice/group, pooled from 7 experiments). Each column was compared with each other column by one-way ANOVA with Tukey’s correction for multiple comparisons as recommended by Prism. Statistics are only shown for relevant comparisons (to simplify visually) but all statistical comparisons and individual values for each mouse are shown in **fig. S7**. ns, not significant, \*\**P* < 0.01, \*\*\**P* < 0.001, \*\*\*\**P* < 0.0001. **(C)** TRAP does not provide late liver-stage control. Schematic for (D). **(D)** Mice were vaccinated in the same groups and imaged by IVIS at 24 and 42 hours as described for (A). However, subgroups of each vaccinated group were treated with anti-CD8 antibodies (αCD8) immediately after imaging at 24-hours. Only imaging data for 42 hours is shown. Mean ± SEM (n = 11 – 13 mice/group, pooled from 3 experiments). Imaging data for both 24 and 42 hours in untreated (no anti-CD8 mAb) mice are also included as a subgroup of total data presented in (B). Indicated pair-wise comparisons were assessed by one-way ANOVA with Sidak correction for multiple comparisons as recommended by Prism. Individual values for each mouse are shown in **fig. S8**. ns, not significant, \*\*\**P* < 0.001, \*\*\*\**P* < 0.0001.

While these findings clearly showed SERA1 was a late-stage target for parasite killing and that TRAP was an early target, they did not directly address whether TRAP-specific T cells could also kill in the late liver-stage. In a set of experiments to address this question, we took advantage of our capacity to deplete CD8^+^ T cells, including the key effector population, liver CD8^+^ Trm cells, by intravenous injection of anti-CD8 mAb (**fig. S4**). To assess killing late in the liver-stage, vaccinated mice were infected with luciferase-expressing parasites and then either left untreated or were injected with anti-CD8 mAb at 24 hours to deplete CD8^+^ T cells, thus preventing parasite killing during the 24 - 42 hour window (**Fig. 4C**). Measurement of luciferase expression showed that CD8^+^ T cell depletion at 24 hours led to increased parasite load in the SERA1-vaccinated group indicating SERA1-specific killing occurred during this 24 - 42 hour period, as expected (**Fig. 4D**). In contrast, CD8^+^ T cell depletion at 24 hours did not affect parasite load in the TRAP-vaccinated group, indicating that TRAP-specific CD8^+^ T cells did not contribute to killing parasite-infected hepatocytes late in the liver stage. As expected, by combining responses to both specificities, maximal parasite killing was achieved, some of which occurred post-24 hours. Together, these findings showed that TRAP-specific Trm cells only significantly kill parasites on the first day of liver-stage infection, while SERA1-specific Trm cells only kill on the second day.

## Discussion

Here we report the discovery of a new PbA liver-stage antigenic T cell target called SERA1. This antigen is expressed late in the liver stage and, as such, only provides an effective target during this temporal window. It is notable that not all liver-stage-expressed proteins can act as targets for killing of infected hepatocytes (*43, 44*). Examples such as GAP50 and 60S ribosomal protein L3 (RPL3) show that despite expression during this stage and their ability to induce CD8^+^ T cell responses, their CD8^+^ T cell effectors are not protective. Knowledge of what enables some liver-stage proteins to act as targets is scant but may relate to whether they can be presented specifically by hepatocytes. While many antigens can be captured and presented by conventional dendritic cells to prime a diverse repertoire of anti-parasite T cells, only a limited fraction of these antigens may be presented by infected hepatocytes. It has been proposed that antigens located within the parasitophorous vacuole do not have access the MHC I-processing machinery and it is only those antigens exported from this vacuole that supply epitopes for presentation (*45*). As such, the location of SERA1 within the membrane of the parasitophorous vacuole (*34*) may be key to its presentation by providing a direct link to the cytosol.

It is less clear how sporozoite proteins like TRAP and CSP access the hepatocyte MHC I processing pathway, since they are associated with the infecting sporozoite, but perhaps cytosolic access is enabled during the invasion process, despite invasion being mediated by vacuole formation. The requirement for cytosolic exposure may also underlie why TRAP only acts as an early target despite evidence for weak mRNA expression by liver stage parasites (*32, 39, 40*). During the hepatocyte-entry phase, sporozoite-associated TRAP may be available for processing onto MHC I molecules, but as suggested by our data, these molecules are turned over rapidly, such that this supply only lasts for a single day. TRAP protein, even if expressed later in the liver-stage, may be much less abundant and restricted to the parasitophorous vacuole, unable to access the cytosolic MHC I machinery.

The demonstrated lack of TRAP-specific killing during the second day of liver-stage infection with PbA aligns well with an earlier observation that, on a per cell basis, Trm cells specific for RPL6, an antigen expressed throughout the liver stage, are more efficient than those specific for TRAP (*29*). This difference would therefore relate to the relative amount of time each specificity has to kill infected cells. While temporally limited presentation of TRAP has a modest impact on killing in mice, extrapolation to humans raises the possibility of a much more detrimental impact. The liver stage for *P. falciparum* extends to about one week but if killing here is also limited to the first 24 hours, then only 14% (1 of 7 days) of the available liver stage killing period will be utilized for TRAP-focused vaccines. This suggests a dramatic gain in efficacy could be achieved by identifying and targeting antigens presented throughout the human liver stage. This finding also brings into question another antigen that is similarly associated with the sporozoite and carried into the hepatocyte, i.e. CSP. For this antigen, mRNA and protein are also known to be expressed at low levels during the liver stage (*46, 47*), but whether its protein products reach the hepatocyte surface on MHC I molecules is unknown. It raises the possibility that CD8^+^ T cell-directed vaccines targeting CSP may also be hampered by a limited period of antigen availability on the infected hepatocyte surface.

With respect to the use of SERA1 as a liver-stage target in human malaria vaccines, this protein originates from a multigene family consisting of five members in PbA and nine members in *P. falciparum* (*48, 49*) and the ortholog of PbA SERA1 is unclear. Each member of the SERA family encodes either a canonical cysteine residue or an atypical serine residue in the catalytic site of its central protease domain. Phylogenetic analyses further categorize SERA genes into four groups (Group I-IV), with all serine-type SERAs (PbA SERA1 and PfSERA1-5) clustering in Group IV and cysteine-type SERA clustering in Group I-III (*48, 49*). As such, PbA SERA1 has five potential orthologs in *P. falciparum*, for which there is limited knowledge of liver-stage expression. A previous study showed that *PfSERA3* is transcribed by *P. falciparum* RAS early after hepatocyte infection (*50*) indicating potential to act as a liver stage target, but clearly more understanding of gene expression in the liver stage is essential to identify the likely ortholog.

Our discovery of TRAP and SERA1 as early and late antigenic targets, respectively, has enabled assessment of the efficacy of combining antigens with different temporal expression windows in a single vaccination. While neither TRAP nor SERA1-induced CD8^+^ T cells were able to fully control PbA infection in all mice, combined vaccination to both antigens induced sterile protection. This was not because we increased the total number of liver Trm cells, the key effector population, because vaccination conditions that yielded similar numbers of Trm cells were still more effective with the combination of antigens. These findings indicate that it is important to understand when during the liver stage a vaccine candidate antigen will be targeted for parasite killing to ensure maximal coverage and parasite clearance. These findings also imply that it is best to target antigens that are expressed throughout the liver stage but that addition of antigens that target defined periods may be beneficial if these periods are not well covered by other vaccine antigens.

## Acknowledgments

We thank the BRF at the Peter Doherty Institute, and the BRU and the Hugh Green Cytometry Centre at the Malaghan Institute of Medical Research, for their technical support. We also thank Melanie Damtsis for her technical assistance.

## Funding

New Zealand Ministry of Business Innovation and Employment contract RTVU1603 (GFP)

New Zealand Health Research Council contract HRC-20/569 (GFP)

New Zealand Health Research Council 14/1003 Malaghan Institute (IFH).

National Health and Medical Research Council of Australia (NHMRC) 1139486 (DFR)

NHMRC 1159272 (SG)

NHMRC 2012701 (LEH, WRH)

NHMRC 1176955 (JAB)

NHMRC 1154457 (WRH)

NHMRC 2016596 (AWP)

## Author contributions

Conceptualization: WRH, IFH, GFP, DFR,

Methodology: YCC, LEH, DFR, CJ, SD, RJA, BJC, DJ, SG, GIM, RBS, TP, RWJS, JAB, IFH, GFP, WRH

Investigation: YCC, LEH, DFR, SD, RBS, RWJS, AC, CJ, SC, DJ

Visualization: YCC, LEH, WRH

Funding acquisition: WRH, LEH, GFP, IFH, SD, DFR, SG

Project administration: WRH, LEH, SD, GP, IFH, GIM, AWP, DFR

Supervision: WRH, LEH, GFP, GIM, AWP

Writing – original draft: YCC, WRH, LEH

Writing – review & editing: YCC, LEH, DFR, SD, RAJ, BJC, RBS, AC, CJ, SC, RWJS, DJ, SG, JAB, AWP, GIM, IFH, GFP, WRH.

## Data and materials availability

All data are available in the main text or the supplementary materials.

