## Supplemental file for "Temporal expression of liver-stage malaria antigens shapes vaccine efficacy"

**The PDF file includes:**

Materials and Methods
Figs. S1 to S9
Tables S1
References (*51-57*)

**Other Supplementary Materials for this manuscript include the following:**

### Supplementary Materials

### Materials and Methods

#### Mice

Mice were either bred in-house at the Peter Doherty Institute [(PDI), The University of Melbourne] or purchased directly from the Animal Resources Centre (Canning Vale, Australia). All mice were maintained at the Bioresource Facility of PDI and experiments performed on animals 6 – 12 weeks of age. For vaccine studies mice used were B6 (Jackson Laboratories), H-2K$^{b-/-}$ mice (*51*) and H-2D$^{b-/-}$ mice (*51*). All animal experiments were in accordance with the Prevention of Cruelty to Animals Act 1986, the Prevention of Cruelty to Animals Regulations 2008, the National Health and Medical Research Council (2013) Australian code for the care and use of animals for scientific purposes, and were approved by the Melbourne Health Research Animal Ethics Committee, University of Melbourne (ethics protocols IDs: 1714302, 1914923, 25361) or the Walter and Eliza Hall Institute of Medical Research Animal Ethics Committee (Approval number 2021.064).

#### Sporozoites and challenges

Animals used for the generation of the sporozoites were 4 – 5-week-old male Swiss Webster mice purchased from Monash Animal Services (Melbourne, Victoria, Australia) and maintained at the School of Botany, The University of Melbourne, Australia. *Anopheles stephensi* mosquitoes (strain STE2/MRA-128 from BEI Resources, The Malaria Research and Reference Reagent Resource Center) were reared as described (*52*). To raise infectious mosquitoes, the *Plasmodium* species used were *P. berghei* ANKA (PbA) wild-type Cl15cy1 (BEI Resources, NIAID, NIH: MRA-871), *PbA-GFP/Luciferase, PbA Mei2*$^{–}$ and *SERA1*$^{–}$ sporozoites. Sporozoites were dissected from mosquito salivary glands (*53*) and resuspended in cold PBS. For challenge experiments, 200, 3000 or 30,000 freshly dissected sporozoites were i.v injected. Blood samples were assessed for parasitemia on days 6, 7, 8, 10 and 12 by flow cytometry after staining with Hoechst 33258 dye (ThermoFisher) for 1 h at 37 °C. An LSR Fortessa (BD Biosciences) with a violet laser (405 nm) was used to excite the dye in infected red blood cells, with percentages of Hoechst-positive cells compared to uninfected controls. Values of > 0.1% were considered positive for parasites, with mice positive for two consecutive days euthanized. Those remaining parasitemia-negative on day 12 were considered protected (*21*).

#### Generation of knockout parasites

The following was performed with the gene region for SERA1 reversed so that the coding sequence reads left to right. 5’ and 3’ homology flanks were amplified from wildtype PbA genomic DNA with the primers CJ616 + CJ617, and CJ618 + CJ619 (Supplementary Table 1), producing amplicons of 822 bp and 805 bp respectively. Both 5’ and 3’ flanks are flush with the ORF for SERA1 and will therefore excise the entire gene sequence. The 5’ flank reverse primer CJ617 and the 3’ flank forward primer CJ618 both featured a 23 base multi-cloning-site complementary tail that was utilized in a further PCR reaction to generate the 5’, 3’ flank fusion fragment. This was cloned into the CRISPR Cas9 plasmid pYC_L2 (a kind gift from Ashley Vaughan) with the restriction enzymes *Hind*III and *Eco*RI. A guide sequence (Guide 5) was identified with

CHOPCHOP and cloned into a KO flank containing plasmid with the enzyme *Esp*3I to generate the SERA1 KO plasmid CJ189 (*54*). 30 µg of pCJ189 was transfected into magnet-purified WT PbA schizont stage parasites as previously described (*42, 55*). In turn, transfected parasites were intravenously injected into Swiss Webster mice (*42*). After 24 hours, mice were given water containing 70 µg/mL pyrimethamine (Sigma Aldrich, USA). When mice surpassed 1% parasitemia, they were euthanized and cardiac bled for cryostocks and genomic DNA extraction. Genotyping was performed by PCR (**table S1**) and clonal parasites generated by limiting dilution as previously described (*55*). *Mei2* KO (*Mei2*⁻) parasites were generated as above except flanks were designed to excise 678 bp upstream and 488 bp downstream of the mei2 coding region. Restriction sites, primers and guides used to generate this KO line are described in table S1.

### Vaccination

Wild-type PbA (40,000 or 50,000) were irradiated with 20,000 rads using a gamma $^{60}$Co source and then i.v. injected into recipient mice (*14*). Alternatively, B6 mice were immunized with three doses of 10,000 freshly isolated PbA *Mei2*⁻ sporozoites at 1-week intervals. For chemoprophylaxis with sporozoites (CPS) vaccinations, sporozoite-infected mice were orally treated with chloroquine (50 mg/kg) starting 4 hours post-infection, with CQ treatments continued for 7 consecutive days. For vaccination with blood stage parasites, a donor B6 mouse was established after intraperitoneal (i.p.) infection with cryopreserved PbA infected red blood cells (iRBC). 4 days later, blood was isolated from the donor mouse via cardiac puncture, measured for parasitemia as described above, and diluted to $10^6$ iRBC in 200 µL for intravenous injection into naïve B6 mice. To cure the blood-stage infection, the infected mice were i.p. treated with CQ (40 mg/L) daily from days 4-8 post-infection, followed by the provision of CQ-infused drinking water (600 mg/L) for another 5 days.

The SERA1 GLP and TRAP GLP vaccines were prepared by oxime conjugation of the N-terminal aminooxy peptides AoAA-FFRK-SKISPDFYNNL and AoAA-FFRK-SALLNVDNL, respectively, to an α-GalCer prodrug with a pendant ketone (**fig. S9**) using methods previously described (*21*). For intravenous administration, compounds were formulated into a tween-histidine-sucrose matrix and reconstituted in distilled water then further diluted in phosphate-buffered saline (PBS) to 0.135 nmol (*56*).

### CD8 enrichment

To enrich $CD8^+$ T cells by negative selection, cells were incubated with $CD8^+$ T cell enrichment cocktail (10 µL per $10^6$ cells) on ice for 30 min. After washing with 10 mL RPMI, 2% FCS, cells were incubated with pre-washed goat anti-rat IgG magnetic beads (1:10 cell: bead ratio) for 20 min on a roller at 4 °C. The tube containing the bead-cell mixture was then placed in a magnetic rack, from which the supernatant containing unbound cells was isolated. A fraction of cells (30 µL) was either used to determine cell count by haemocytometer or to check the purity of CD8 T cells by flow cytometry.

### ELISpot

96-well ELISpot plates were washed five times with PBS, coated with 3 µg/mL IFN-γ capture Ab diluted in 80 µL PBS, and then incubated overnight at 4 °C. The coated plates were washed with

200 μl PBS five times before blocking with 100 μL of RPMI, 10% FCS for 30 min at room temperature. Splenocytes isolated from pooled immunized mice were enriched for $CD8^+$ T cells, as described above. Next, 250,000 enriched $CD8^+$ T cells were plated and stimulated in the presence or absence of 5 μM peptide for 18-20 hours at 37 °C and 5% $CO_2$ in a humidified incubator. After incubation, the plates were washed 5 times with 200 μL of PBS then incubated with 1 μg/mL biotin-conjugated detection antibody diluted in PBS, 0.5% FCS for 2 hours. Plates were again washed 5 times with 200 μL of PBS followed by 1 hour incubation with 100 μL of streptavidin-HRP diluted in PBS, 0.5% FCS, and finally with 100 μL of TMB substrate for 20 min. After the final washing, plates were dried overnight and IFN-γ spots were enumerated by an AID ELISpot reader (Advanced Imaging Devices GmbH, Strassberg, Germany).

**Tissue preparation and flow cytometry**

Livers were collected into tubes containing RPMI, 2% FCS and 10 U heparin. Each liver was teased through a 70 μm cell strainer, washed with RPMI, and the cell suspension resuspended in 30 mL of 35% Percoll (GE Health care) before centrifugation at 500 x g for 20 min at 20-22 °C, with no brake. The cell pellet was incubated in 5 mL RBC lysis solution for 5 min at 20-22 °C, before being washed with 30 mL RPMI and resuspended in 1 mL of FACS buffer (PBS + 5% w/v BSA + 5 mM EDTA) for flow cytometry. One quarter of the cell suspension was used for antibody staining. Spleen lymphocytes were isolated by teasing tissue through a 70 μm strainer, washing in RPMI, 2% FCS, incubating cells in RBC lysis solution for 1-2 min at 20-22 °C, washing again and resuspending in FACS buffer. Approximately $1/25^{th}$ of final volume was used for antibody staining. Lymphocytes were stained with tetramers for 1 hour at 20-22 °C before staining with surface antibodies. Samples were run on an LSRFortessa (BD Biosciences) and data were analysed using Flowjo X software (TreeStar).

**Precursor numbers**

Single-cell suspensions isolated from the spleen and major peripheral lymph nodes (axillary, branchial, mesenteric, cervical and inguinal) of naïve B6 mice were incubated with Fc blocking cocktail (24G2) and 100 nM dasatinib at 37 °C for 30 min. PE-conjugated tetramer (1/200 dilution) was diluted in 100 μL of FACS buffer and incubated at room temperature in the dark for 1h. After washing with cold FACS buffer, tetramer-labelled cells were then incubated with 50 μL of anti-PE microbeads at 4 °C for 20 min. Cells were washed with FACS buffer and a positive enrichment was performed using an LS column (Miltenyi Biotech) in a magnetic rack. The enriched samples were stained with monoclonal antibodies specific for TCRβ, CD8, CD19, F4/80, NK1.1, CD44 and CD62L for 30 min on ice before the whole sample was acquired on a flow cytometer.

**IVIS imaging**

B6 mice were challenged i.v. with 30,000 luciferase expressing sporozoites and parasite burden assessed at 24 and 42 hours using whole body imaging (IVIS, Xenogen, USA). The luciferase substrate, XenoLight D-luciferin potassium salt (Perkin Elmer), was dissolved in phosphate-buffered saline at a concentration of 5 mg/mL. Mice were anesthetized in an oxygen-rich induction chamber with 3.5% isoflurane and shaved. Whole body imaging was performed 10 min after i.p. injection with 0.05 mg/g body weight. Bioluminescence imaging was acquired with 21.7 cm field of view for whole body imaging, with a medium binning factor, and exposure time of 5 min. For bioluminescence quantification, regions of interest (ROI) were drawn by using the software Living Image 3.0 and average radiance (p/s/cm$^2$/sr) was determined. In some instances, mice were depleted of CD8$^+$ T cells by treatment with 100 μg of anti-CD8 (2.43 clone) i.v. at 24 hours (*14*). CD8$^+$ T cells were depleted within 2 hours (**fig. S4b)**.

To normalize values from each experiment, the relative radiance for each mouse was calculated as a percentage of the average radiance of the unvaccinated (naïve) group at 24 hours post-sporozoite challenge.

**Statistics**

Treatments were assigned to separately caged litter groups. Data analysed included outliers unless explained by technical error. Data collection and analysis were not performed blind to the conditions of the experiments. Data was assumed to be normal, but this was not formally tested because the sample sizes used were not well powered for normality testing. The statistical tests performed on the data are indicated in the figure legends, along with sample size. Individual P values are provided in Figures as follows: *$P < 0.05$, **$P < 0.01$, ***$P < 0.001$, ****$P < 0.0001$. Statistical analysis and figures were generated using GraphPad Prism 9.

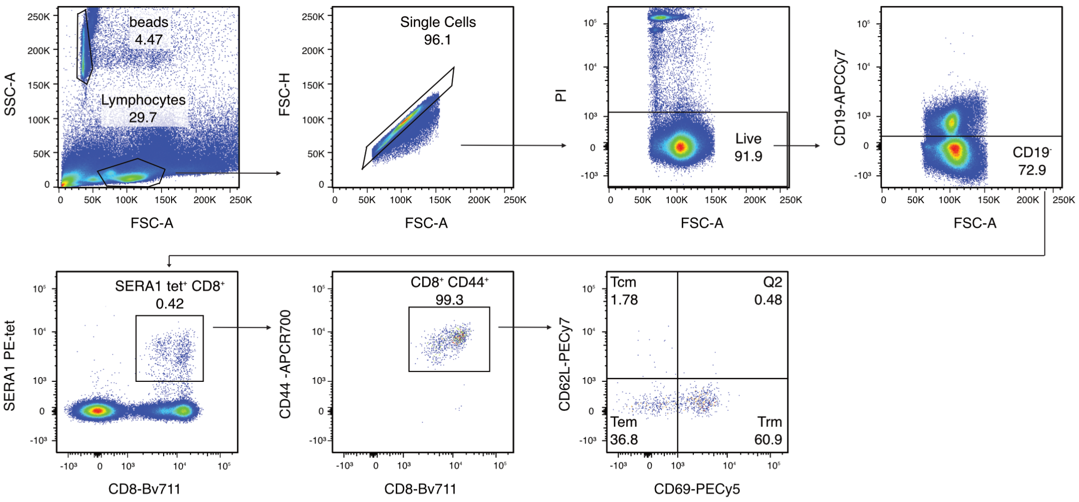


**Fig. S1. Gating strategy for analysis of endogenous SERA1-specific memory CD8$^+$ T cells responding to vaccination.**

All cells were gated on using FSC-A and SSC-A profiles. Doublets were removed from the analysis using FSC-A vs FSC-H and dead cells were excluded by eliminating cells positive for Propidium Iodide (PI). From the live lymphocytes, CD19$^-$ cells were selected, followed by tet$^+$ CD8$^+$ cells and then CD44$^+$ cells, before delineating memory subsets into tissue-resident memory (Trm; CD69$^+$ CD62L$^{low}$), effector memory (Tem; CD69$^-$ CD62L$^{low}$) and central memory (Tcm; CD69$^-$ CD62L$^+$) cells using antibodies against CD69 and CD62L.

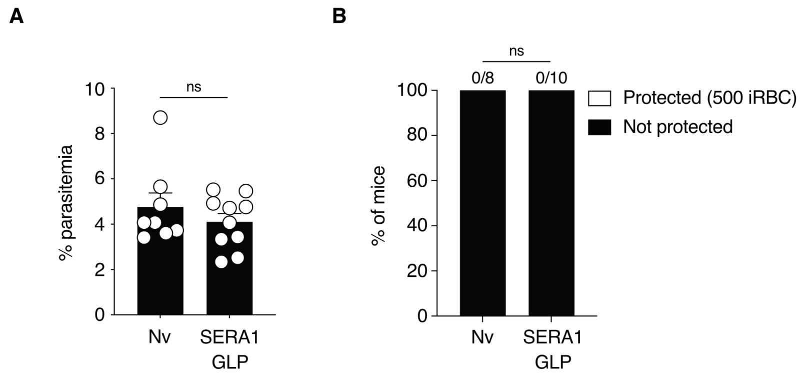


**Fig. S2. Vaccination of B6 mice with SERA1 GLP does not protect against blood-stage malaria challenge.**
B6 mice were immunized intravenously with SERA1 GLP, and 33-42 days later, challenged intravenously with 500 PbA iRBC and measured for parasitemia up to day 7. **(A)** Day 7 parasitemia. **(B)** The proportion of mice that succumbed to malaria infection (black bar) or protected against iRBC challenge (white bar). Numbers above bars indicate the number of mice that were protected against iRBC challenge over the total number of mice challenged. Data are pooled from 2 independent experiments (n = 8 – 10 mice/group). Data in (A) were log-transformed and compared using an unpaired two-tailed *t*-test. Data in (B) was compared using a two-sided Fisher's exact test. ns, not significant.

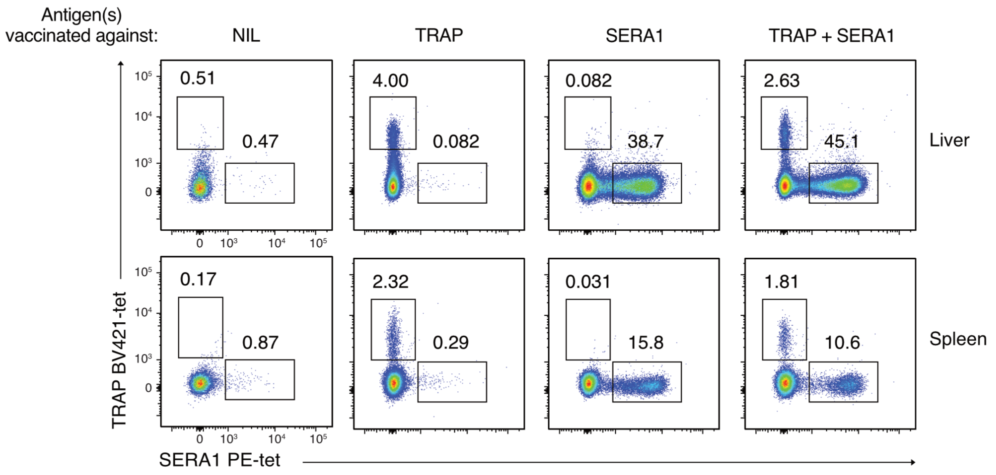


**Fig. S3. Memory CD8$^+$ T cell responses after GLP vaccination.**
Representative FACS plots of memory CD8$^+$ T cells specific for TRAP or SERA1 in the liver (top row) and spleen (bottom row) after vaccination with single or combined antigens as described in Fig. 3A.

**A**

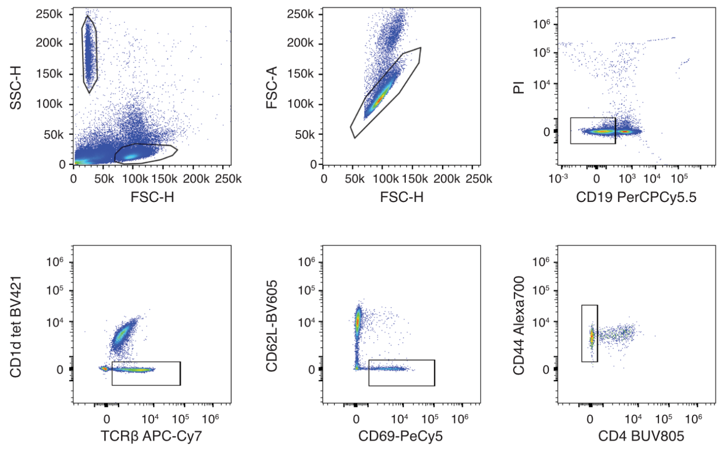


**B**

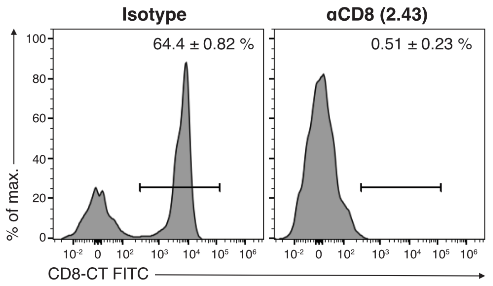


**Fig. S4. In vivo depletion of liver CD8$^+$ Trm cells.**

B6 mice were injected i.v. with 100 µg of anti-CD8 (2.43 clone) or left untreated. **(A)** Livers were harvested 2 hours later and stained with a CD1d tetramer and a CD19 antibody to exclude B and NKT cells from the analysis. Conventional T cells were gated on using a TCRβ$^+$ antibody then Trm cells were identified by gating on CD69$^+$ CD62L$^{low}$ cells. CD4$^+$ T cells were excluded from the analysis and CD44$^+$ cells selected. **(B)** CD8$^+$ Trm cells were identified by staining with CD8-CT antibody, a clone previously shown to retain binding capacity after treatment with the *in vivo* depleting antibody 2.43 (*57*). Histograms are representative of one experiment containing 3 mice.

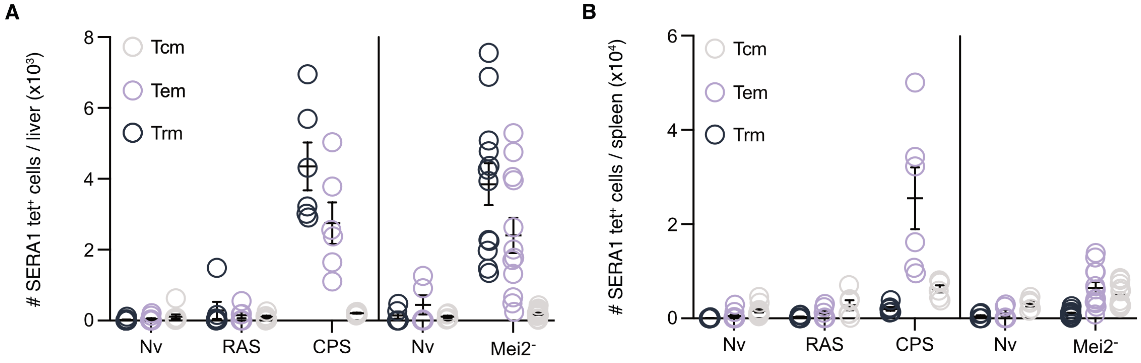


**Fig. S5. Individual data points for Fig. 1E and F.**
(**A**). Individual data points for Fig. 1E. (**B**). Individual data points for Fig. 1F

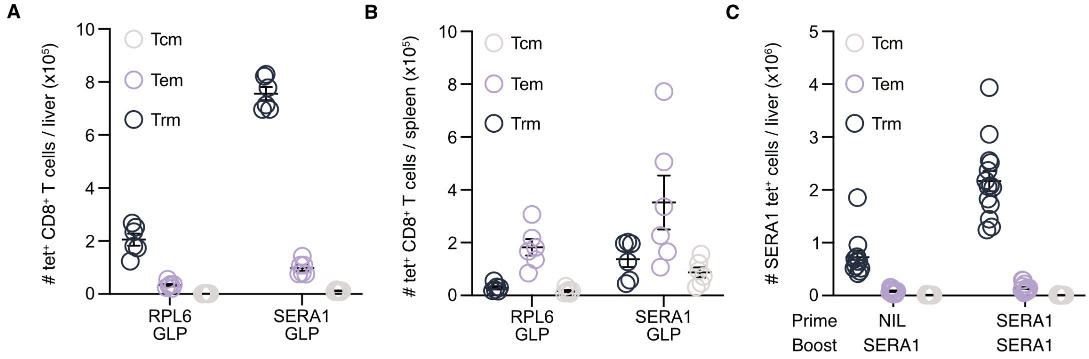


**Fig. S6. Individual data points for Fig. 2C, D and J.**
(**A**). Individual data points for Fig. 2C. (**B**). Individual data points for Fig. 2D. (**C**). Individual data points for Fig. 2D.

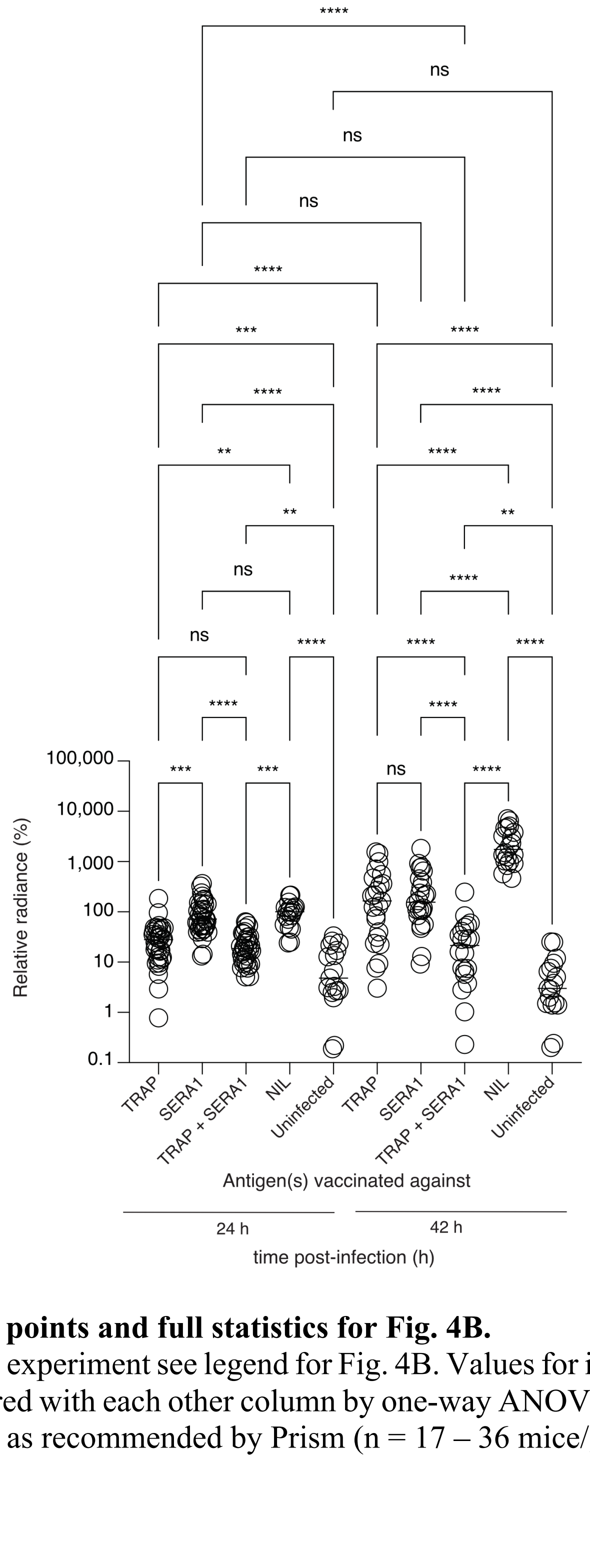


**Fig. S7. Individual data points and full statistics for Fig. 4B.**
For information about the experiment see legend for Fig. 4B. Values for individual mice are shown. Each column was compared with each other column by one-way ANOVA with Tukey's correction for multiple comparisons as recommended by Prism (n = 17 – 36 mice/group).

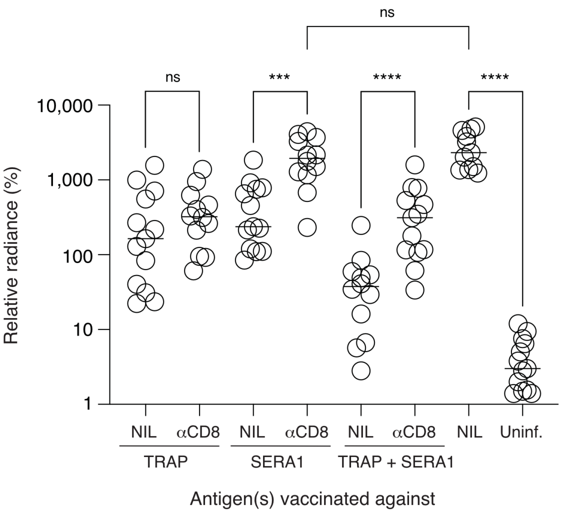


**Fig. S8. Individual data points and full statistics for Fig. 4D.**
For information about the experiment see legend for Fig. 4D. Indicated pair-wise comparisons were assessed by one-way ANOVA with Sidak correction for multiple comparisons as recommended by Prism (n = 11 – 13 mice/group).

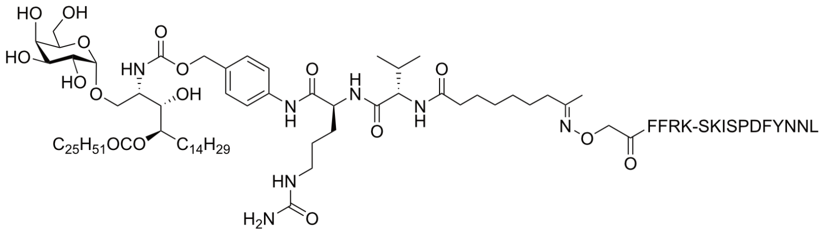


GLP-SERA1

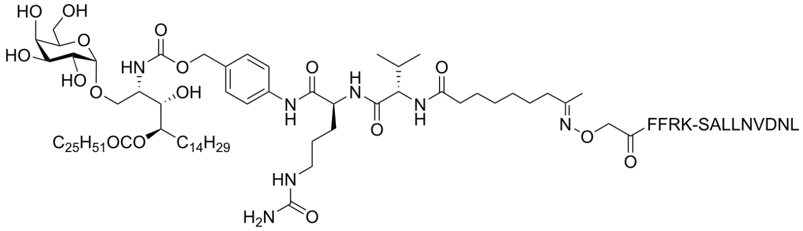


GLP-TRAP

**Fig. S9. Chemical structure of the SERA1 GLP and TRAP GLP vaccines.**

**Table S1. Restriction sites, primers and guides used for PbA knockout line generation**

| Name | Gene | F/R | Sequence | Purpose |
|---|---|---|---|---|
| PbSERA1 Guide 5-F | SERA1 | F | tattATTAGTGGAACTGGACACGA | Guide for PbSERA1 |
| PbSERA1 Guide 5-R | SERA1 | R | aaacTCGTGTCCAGTTCCACTAAT | Guide for PbSERA1 |
| CJ616 | SERA1 | F | ATATAAGCTTGTTATGTTAGTGTATCATTTATGTATTGC | 5' flank forward with HindIII |
| CJ617 | SERA1 | R | CCAACCTAGGTATAGGCGCGCCTTTTTACAAGTATAATATTAAACTGGAACTTG | 5' reverse with linker |
| CJ618 | SERA1 | F | AGGCGCGCCTATACCTAGGTTGGTGATTTTAAGAAATGCTTATGAAATTTATG | 3' forward with linker |
| CJ619 | SERA1 | R | atatGAATTCATATGCAAATAATACTAATACGTCATTATAATACC | 3' reverse with EcoRI |
| CJ620 | SERA1 | F | CCTTAAGAATTTTTGAAAGAATTAATCATCC | Outside 5' flank for genotyping |
| CJ621 | SERA1 | R | CTTACATAATATAAAAAGAATGGAAAGACG | Inside 5' wt |
| CJ622 | SERA1 | F | CTAAATTGTACAATTCAGACGATTGC | Inside 3' wt |
| CJ623 | SERA1 | R | GAATAAAAGCATATAATACATATTGTATGCC | Outside 3' flank for genotyping |
| CJ614 | - | R | CCAACCTAGGTATAGGCGC | Inside in linker for KO genotyping |
| CJ567 | - | F | GCGCCTATACCTAGGTTGG | Inside linker to detect KO |
| CJ472 | mei2 | F | TATAGGTACCGTATGGGAACATGCATATTATGTGG | Forward to amplify the 5' flank of mei2 ko primer KpnI |
| CJ473 | mei2 | R | CCAACCTAGGTATAGGCGCGCCTACAAAAGGAATATGGGGAATACACC | Reverse for 5' Flank with linker including PAM site |
| CJ474 | mei2 | F | AGGCGCGCCTATACCTAGGTTGGGGATGTTTATAAATAAAATAGTGTAAAATTCG | 3' Flank fwd with PAM linker |
| CJ475 | mei2 | R | atatGAATTCCCTGTTTAGGTTTATTTTGTCATTTCAC | 3' flank rev with EcoRI |
| CJ562 | mei2 | F | GGCATGATGCCGAATGCC | Outside 5' flank for genotyping |

| | | | | |
|---|---|---|---|---|
| CJ563 | mei2 | R | CATGTGTACACACGATCATATGG | Inside excised region downstream of 5' flank. Absent in KO |
| CJ564 | mei2 | F | GATGACCCGAGTTGTGAAGG | Inside excised region upstream of 3' flank. Absent in KO |
| CJ565 | mei2 | R | GAATTTTGAATATATGATCATAT CGCACG | Outside 3' flank for genotyping |
| CJ566 | mei2 | R | CTAGGTATAGGCGCGCC | Inside linker to detect KO |
| CJ567 | mei2 | F | GCGCCTATACCTAGGTTGG | Inside linker to detect KO |
| CJ595 | mei2 | F | GCCTATACCTAGGTTGGGG | Alternative fwd in linker region of mei2KO for KO genotyping |
| Mei2 KO guide 28 | mei2 | F | tattCATGTGTACACACGATCATA | Guide for mei2 |
| Mei2 KO guide 28 | mei2 | R | aaacTATGATCGTGTGTACACATG | Guide for mei2 |
| Mei2 KO Guide 3 | mei2 | F | tattAATGATGACCCGAGTTGTGA | Guide for mei2 |
| Mei2 KO Guide 3 | mei2 | R | aaacTCACAACTCGGGTCATCATT | Guide for mei2 |
